# Estimating the vertical ionization potential of single-stranded DNA molecules

**DOI:** 10.1101/2023.02.27.530325

**Authors:** Marianne Rooman, Fabrizio Pucci

**Affiliations:** Computational Biology and Bioinformatics, Université Libre de Bruxelles, 1050 Brussels, Belgium; Interuniversity Institute of Bioinformatics in Brussels, 1050 Brussels, Belgium

## Abstract

The electronic properties of DNA molecules, defined by the sequence-dependent ionization potentials of nucleobases, enable long-range charge transport along the DNA stacks. This has been linked to a range of key physiological processes in the cells and to the triggering of nucleobase substitutions, some of which may cause diseases. To gain molecular-level understanding of the sequence dependence of these phenomena, we estimated the vertical ionization potential (vIP) of all possible nucleobase stacks in B-conformation, containing one to four Gua, Ade, Thy, Cyt or methylated Cyt. To do this, we used quantum chemistry calculations and more precisely the second-order Møller-Plesset perturbation theory (MP2) and three double-hybrid density functional theory (DFT) methods, combined with several basis sets for describing atomic orbitals. The calculated vIP of single nucleobases were compared to experimental data and those of nucleobase pairs, triplets and quadruplets, to observed mutability frequencies in the human genome, reported to be correlated with vIP values. This comparison selected MP2 with the 6-31G* basis set as the best of the tested calculation levels. These results were exploited to set up a recursive model, called vIPer, which estimates the vIP of all possible single-stranded DNA sequences of any length based on the calculated vIPs of overlapping quadruplets. vIPer’s vIP values correlate well with oxidation potentials measured by cyclic voltammetry and activities obtained through photoinduced DNA cleavage experiments, further validating our approach. vIPer is freely available on the github.com/3BioCompBio/vIPer repository.

## Introduction

DNA molecules are not only the repository for genetic information of all living organisms, but they also have unique electronic properties. Indeed, they can act as molecular wires formed by the *π*-stacking of the aromatic moieties of the nucleobases, along which long-range charge transport takes place.^1,2^ Such charge transport even occurs in cells, where exposure to high-energy radiations or to reactive oxygen species generated as by-products of the cellular metabolism can lead to DNA ionization through the creation of electron holes.^3,4^ These holes have been shown to migrate along the nucleobase stack until they remain localized in a region of low ionization potential, where they basically have two consequences: they are repaired by specific enzymes^5^ or trigger single base substitutions.^6^

Recently, several studies have emphasized the link between DNA mutability and the vertical ionization potential (vIP) of the DNA sequence where the mutation occurs. It has indeed been shown that the base substitution rates depend on the DNA sequence context,^7,8^ and that the lower the vIP of the motif formed by a substituted nucleobase and its adjacent DNA sequence, the more likely the substitution, whether for somatic cancer mutations or germline mutations.^6,9,10^ The existence of a link of vIP and mutability with pathogenicity has been suggested and discussed. ^6,7,11^

The various effects and possible roles of electron holes migrating along cellular DNA are not yet fully elucidated. Too many holes are obviously damaging to the cell if they are not repaired in time; for example, deficiency in repair enzymes are known to cause diseases such as cancer or neurodegeneration.^12–14^ But a limited amount of electron holes appears to have a physiological role, as supported by the precise regulation of the amount of reactive oxygen species, which is, e.g., higher during cell differentiation.^15^ Moreover, charge transfer seems to be used by the cell as a signaling mechanism, e.g., to cooperatively signal damage to DNA repair proteins.^5,16^ The biological role of charge transfer is also supported by the recurrent occurrence of specific DNA conformations consistent with charge migration at the interface with specific proteins, ligands and metal ions.^17^

In this fascinating biological context, it is of utmost importance to have an accurate estimation of the ionization potential of nucleobase *π*-stacks. The four nucleobases that make up DNA have different propensities to be ionized, with the consequence that all possible nucleobase motifs have also different vIP values. The vIPs of the individual nucleobases have been measured experimentally,^18^ but not those of sequences of nucleobase *π*-stacks. To estimate the vIPs of nucleobase stacks, quantum chemistry calculations have been performed using Hartree-Fock (HF)^19–21^ and M06-2X, a simple hybrid density functional theory (DFT).^22,23^ However, *π*-*π* stacking interactions involve strong electron delocalization and thus large dispersion contributions,^24,25^ which are not represented well enough in these theories^26^. These calculations nevertheless demonstrated the high dependence of vIP values on DNA sequence, but also on DNA conformations. ^23^ Other ways of estimating the vIPs of nucleobase stacks involves using molecular dynamics simulations in conjunction with various levels of quantum chemistry calculations.^27–29^ However, these methods are too computer time consuming to obtain the vIPs of all possible nucleobase stacks.

To fill this gap, we present here a user-friendly software called vIPer that estimates the vIP of B-form single-stranded DNA stretches of any length based on quantum chemistry calculations on *π*-stacks formed of four successive nucleobases combined with mathematical modeling techniques. Since cytosine methylation has crucial functions within cells and is involved in gene expression, development and cancer,^30^ we considered 5-methylcytosine (5mC) in addition to the four basic DNA nucleobases Ade, Cyt, Gua and Thy.

## Methods

### Quantum chemistry calculations

Since dispersive forces have a huge importance in DNA base stacking, ^24,25^ we used levels of calculation known to account for these forces in a satisfactory manner - one ab initio and three DFT methods.

The ab initio method we used is second-order Møller-Plesset perturbation theory (MP2).^31^ Although MP2 has been shown to systematically overestimate *π*-stacking energy,^32^ this effect is rather weak and not very problematic in the present context as we focus on ionization potentials and thus on energy differences, which we moreover compare between different DNA motifs. MP2 also has the advantage of being an ab initio method that, while considering electron correlations, is relatively inexpensive in terms of computation.

We also used three double-hybrid methods that combine exact HF exchange with MP2-like correlation to DFT, i.e. B2-PLYP,^33^ mPW2-PLYP^34^ and PBE0-DH.^35^ Note that the amount of exact exchange is much bigger in these methods than in the simple hybrids and dispersion forces are more accurately computed. The computational cost of this class of DFT methods is similar to that of MP2.

In addition, we tested several basis sets for atomic orbitals. First we considered the split-valence basis sets 6-31G* and 6-31G** with polarization functions on heavy atoms, and on heavy and hydrogen atoms, respectively.^36,37^ In these basis sets, the exponent *α_d_* of the Gaussian d-polarization function on the heavy atoms is equal to 0.8. We also considered two modified basis sets with *α_d_*=0.2, which we denote here 6-31G^(.2)^ and 6-31G^(.2)^*. The *α_d_*-exponent modulates the spatial extension of the wave function, with lower values corresponding to more diffuse orbitals. The value of 0.2 has been determined to minimize the MP2 energy of a stair motif involving two stacked Gua bases forming a hydrogen bond and a cation-*π* interaction with an Arg residue,^38^ a common motif found at protein-DNA interfaces.^39^ Note that *α_d_*=0.2 is close to the value of 0.25 used by other authors to estimate the stacking energies between aromatic systems.^24^

All quantum chemistry calculations were performed in gas phase and with the Gaussian 16 suite.^40^

### Single nucleobase and nucleobase stack geometries

We considered the four nucleobases Ade (A), Cyt (C), Gua (G) and Thy (T) whose combinations form DNA molecules. In addition, we considered 5-methylcytosine (5mC or M) that is the common methylated form of cytosine. In these five nucleobases, the sugar cycle and phosphate group were omitted, and the glycosidic bond was replaced by a hydrogen atom.

The initial geometries of the single nucleobases were taken from the software package x3DNA-DSSR.^41^ These geometries were optimized using the different levels of theory and basis sets described in the previous subsection.

In a second stage, we considered single-stranded nucleobase stacks in standard B conformation. The geometries of the stacks were designed using x3DNA-DSSR. This software proposes two generic B-DNA conformations that both have a twist of 36.0°. The first, noted here B1 (B55 in x3DNA-DSSR) has a rise of 3.39 Å,^42^ and the second, noted B2 (B4 in x3DNA-DSSR), a rise of 3.375 Å.^43^ We then replaced each of the nucleobases forming the stacks with the same nucleobases but with optimized geometry. This was done by superimposing the optimized bases onto the non-optimized ones by minimizing the root mean square deviation of atomic positions using the U3BEST algorithm. ^44^ Examples of such stacks are shown in Fig. 1.

**Figure 1:**
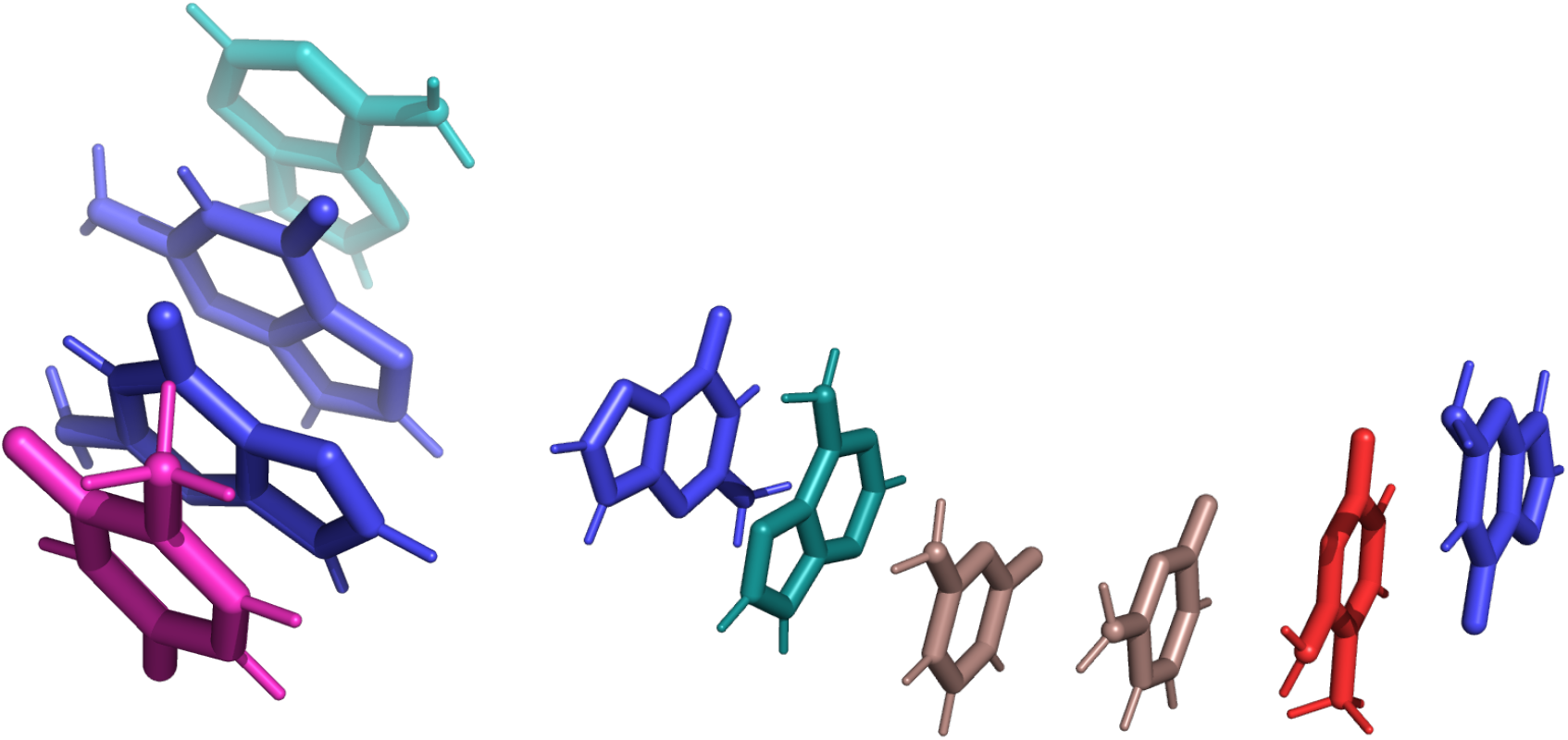
Single-stranded B1 stacking geometry of a TGGA nucleobase quadruplet and a GACCMG sextuplet. Gua are in blue, Ade in teal, Thy in pink, Cyt in darksalmon and 5mC in red.

### Vertical ionization potential (vIP)

We considered two species of each molecule: a neutral species *S* and a radical cationic species *S*^•+^ with one missing electron. The vIP is defined as the difference in energy between these two species, both considered to adopt the optimal conformation *C* of the neutral species, thus ignoring possible changes in molecular geometry that may result from ionization:

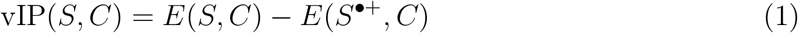

The energy calculations were performed using the levels of theory and basis sets described in the previous subsections. For cationic species, they were performed with restricted open-shell procedures to prevent spin contamination problems.

## Results

### vIP of single nucleobases

As a first stage, we computed the vIP of the four nucleobases Gua, Ade, Cyt and Thy at MP2, B2-PLYP, mPW2-PLYP and PBE0-DH levels of theory with 6-31G*, 6-31G**, 6-31G^(.2)^ and 6-31G^(.2)^* basis sets (see Methods), and compared them with experimentally measured vIP values.^18^ The results are given in Table 1 and Supplementary Table S1.

**Table 1:** Calculated and experimental vIP values (in eV) of single nucleobases. The root mean square error (RMSE; in eV) and the Pearson correlation coefficient (*r*) with the associated *p*-values are between calculated and experimental^18^ vIP values.

| Level of theory | MP2 |  | B2-PLYP |  | Experi<br>mental<sup>18</sup> |
| --- | --- | --- | --- | --- | --- |
| Basis set | 6-31G* | 6-31G<sup>(.2)</sup> | 6-31G* | 6-31G<sup>(.2)</sup> |  |
| Gua | 7.86 | 7.99 | 7.71 | 7.83 | 8.24 |
| Ade | 8.30 | 8.37 | 8.04 | 8.17 | 8.44 |
| Cyt | 8.56 | 8.56 | 8.43 | 8.54 | 8.94 |
| Thy | 8.77 | 8.88 | 8.72 | 8.83 | 9.14 |
| RMSE | 0.33 | 0.26 | 0.47 | 0.35 |  |
| $r$ | 0.96 | 0.96 | 0.99 | 0.99 | |
| $p$-value | 0.04 | 0.04 | 0.009 | 0.01 | |

| Level of theory | mPW2-PLYP |  | PBE0-DH |  |  |
| --- | --- | --- | --- | --- | --- |
| Basis set | 6-31G* | 6-31G<sup>(.2)</sup> | 6-31G* | 6-31G<sup>(.2)</sup> |  |
| Gua | 7.75 | 7.86 | 7.94 | 8.00 |  |
| Ade | 8.08 | 8.20 | 8.27 | 8.35 |  |
| Cyt | 8.49 | 8.59 | 8.73 | 8.79 |  |
| Thy | 8.77 | 8.88 | 8.98 | 9.02 |  |
| RMSE | 0.42 | 0.31 | 0.22 | 0.16 |  |
| $r$ | 0.99 | 0.99 | 1.00 | 0.99 | |
| $p$-value | 0.008 | 0.01 | 0.005 | 0.007 | |

We observe that, when considering the nucleobases individually, all the levels of theory and basis sets that we tested give very accurate results, with a slight preference for double hybrid DFT methods, and in particular PBE0-DH/6-31G* that reaches *r* =1 and PBE0-DH/6-31G^(.2)^ that yields RMSE=0.16 eV. Very low difference is observed between 6-31G* and 6-31G**, and between 6-31G^(.2)^ and 6-31G^(.2)^*. Note, however, that there is an uncertainty of about ±0.03 eV on the experimental values,^18^ which means that the observed differences in RMSE and *r* must be considered with caution.

### vIP of nucleobase pairs

We calculated the vIP values of all 16 pairs of successive nucleobases containing Gua, Ade, Thy and Cyt in B1 conformation, using the four levels of theory and the four basis sets considered here (see Methods). The results are given in Table 2 and Supplementary Table S2. We considered only single-stranded DNA molecules, as electron holes are known to generally migrate along a single strand and to only rarely jump onto the complementary strand.^45^

**Table 2:** Calculated vIP values (in eV) of pairs of successive nucleobases in B1 conformation. The Pearson correlation coefficient *r* and associated *p*-values are computed between the calculated vIP and log 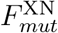, the logarithm of the mutation frequency.

| Level of theory | MP2 |  | B2-PLYP |  | mPW2-PLYP |  | PBE0-DH |  |
| --- | --- | --- | --- | --- | --- | --- | --- | --- |
| Basis set | 6-31G* | 6-31G<sup>(.2)</sup> | 6-31G* | 6-31G<sup>(.2)</sup> | 6-31G* | 6-31G<sup>(.2)</sup> | 6-31G* | 6-31G<sup>(.2)</sup> |
| GG | 7.47 | 7.61 | 7.27 | 7.42 | 7.32 | 7.45 | 7.47 | 7.56 |
| GA | 7.73 | 7.78 | 7.51 | 7.64 | 7.56 | 7.67 | 7.69 | 7.80 |
| GC | 7.76 | 7.81 | 7.58 | 7.69 | 7.61 | 7.72 | 7.78 | 7.86 |
| GT | 7.74 | 7.86 | 7.57 | 7.69 | 7.61 | 7.72 | 7.79 | 7.85 |
| AG | 7.57 | 7.71 | 7.26 | 7.48 | 7.34 | 7.56 | 7.52 | 7.62 |
| AA | 7.99 | 7.77 | 7.50 | 7.67 | 7.58 | 7.74 | 7.78 | 7.89 |
| AC | 8.25 | 8.25 | 7.96 | 8.08 | 8.00 | 8.12 | 8.17 | 8.26 |
| AT | 8.17 | 8.24 | 7.86 | 8.01 | 7.92 | 8.06 | 8.09 | 8.18 |
| CG | 7.41 | 7.55 | 7.23 | 7.38 | 7.28 | 7.40 | 7.44 | 7.53 |
| CA | 7.85 | 7.94 | 7.56 | 7.70 | 7.60 | 7.73 | 7.78 | 7.87 |
| CC | 8.07 | 8.04 | 7.88 | 8.00 | 7.99 | 8.10 | 8.15 | 8.24 |
| CT | 8.34 | 8.42 | 7.89 | 8.02 | 7.98 | 8.11 | 8.24 | 8.32 |
| TG | 7.73 | 7.85 | 7.52 | 7.67 | 7.58 | 7.70 | 7.75 | 7.83 |
| TA | 8.16 | 8.21 | 7.85 | 7.98 | 7.90 | 8.02 | 8.08 | 8.16 |
| TC | 8.19 | 8.16 | 8.14 | 8.27 | 8.24 | 8.36 | 8.46 | 8.54 |
| TT | 8.11 | 8.18 | 8.13 | 8.25 | 8.22 | 8.34 | 8.48 | 8.54 |
| $r$ | -0.80 | -0.70 | -0.73 | -0.75 | -0.72 | -0.74 | -0.71 | -0.72 |
| $p$-value | $2 \times 10^{-4}$ | $2 \times 10^{-3}$ | $1 \times 10^{-3}$ | $8 \times 10^{-4}$ | $2 \times 10^{-3}$ | $9 \times 10^{-4}$ | $2 \times 10^{-3}$ | $2 \times 10^{-3}$ |

Here, we do not have experimental vIP values to compare with the calculated vIPs, and we have thus no means to select the best level of theory and basis set. A possibility would be to compare our calculated vIPs with those obtained with gold-standard ab initio quantum chemistry methods such as CCSD(T),^46^ but these are much too computer intensive to be applied to calculate the vIPs of all possible nucleobase pairs, triplets and quadruplets.

Another possibility is to correlate the calculated vIP values with mutability data. Indeed, we showed earlier^6^ that there is a statistically significant anticorrelation between the frequency of single base substitutions in human genomes and the vIP of the motif consisting of the substituted base and its flanking DNA sequence. In a nutshell, the lower the vIP value, the more probable the substitution. We chose this point of view here and stated that the quantum chemistry method which yields the best anticorrelation is also the most reliable method for calculating the vIP of the considered molecules.

The rationale for supposing a relationship between vIP and mutability originates from the effect of oxidative stress. In the cells, this stress is due to physical or chemical agents, such as exposure to ionizing radiation, long-wave ultraviolet light, and reactive oxygen species. It often causes the extraction of electrons from the DNA molecules. The electron holes so created then migrate along the DNA stack towards regions of low vIP, where they are either repaired by specific enzymes or are likely to mutate.^3–6^

To get mutability data, we used the ClinVar database,^47^ which contains essentially germline variants in human genomes and their associated phenotypes. We focused here on synonymous variants in exon regions. The reason of this choice is that missense mutations cause amino acid substitutions in the encoded proteins, of which some are likely to affect protein stability or function too strongly, so that they do not get fixed. The anticorrelation of the vIP values with synonymous variant frequencies are therefore expected to be better than when considering missense mutations, which is indeed what is observed. ^6^

We thus computed from the ClinVar database the frequencies of all possible base pair motifs with the first base mutated, referred to as 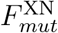, where X denotes the mutated position and N any base. The 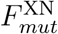 values and details about their calculation can be found in a previous article. ^6^ We then correlated the vIP values for all base pairs with the logarithm of their mutation frequency (log 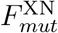). The Pearson correlation coefficient *r* is given in Table 2 and Supplementary Table S2. We first observe that the correlation is negative, as expected, as lower vIP values attract electron holes, which triggers mutation mechanisms.

Moreover, we see that *r* is by far the most negative for MP2 level of theory compared to the considered double-hybrid DFT methods. The result here is thus different than for individual nucleobases, where the DFT methods appeared slightly superior. The reason is that, in pairs of stacked nucleobases, *π*-orbitals overlap and electron delocalization is an important contribution to stability, whereas it is not the case for individual bases. We conclude that for interactions involving aromatic stacking between nucleobases, MP2 appears as a better level of theory than the tested double-hybrid DFT methods.

Moreover, we observe that the basis sets 6-31G^(.2)^ and 6-31G^(.2)^*, which represent more diffuse orbitals than the usual 6-31G* and 6-31G** sets (see Methods), lead to slightly better correlation coefficients when associated with DFT calculations, but to worse correlations when associated with MP2. This contrasts with the results obtained for interaction energies at MP2 level.^24,38^ Furthermore, adding polarization on hydrogen atoms, i.e. considering the 6-31G** set rather than 6-31G*, and 6-31G^(.2)^* rather than 6-31G^(.2)^, does not improve the results on the average. As the inclusion of these polarization functions is computationally expensive, we choose not to consider them in what follows.

We also calculated the vIP values of all the nucleobase pairs in B2 conformation at MP2 level of theory using the four different basis sets considered (Supplementary Table S3). Clearly, the vIP values calculated for nucleobase stretches adopting this type of B-conformation are less well correlated with the mutation frequency than those in B1 conformation. This corroborates the high sensitivity of vIP values to DNA conformation previously observed, ^23^ and explains the lower vIP-frequency correlations that we found in a previous study. ^6^ This result suggests that the B1 conformation is closer to the average DNA conformation adopted in the cell than the B2 conformation.

In summary, on the basis of the correlation between vIP and mutability values, we conclude that the best of the tested quantum chemistry methods for DNA stacks is MP2/6-31G*, and that the most realistic of the tested single-stranded DNA stack conformations is B1. We thus limit ourselves to this level of theory and this DNA conformation in what follows.

### vIP of triplets and quadruplets

We used the MP2/6-31G* quantum chemistry theory to estimate the vIP of all 64 base triplet motifs and all 256 base quadruplet motifs. The results are shown in Table 3 and Supplementary Table S4. The range of vIP values increases with the number of bases in the motif and the average vIP value decreases, as shown in Fig. 2.

**Figure 2:**
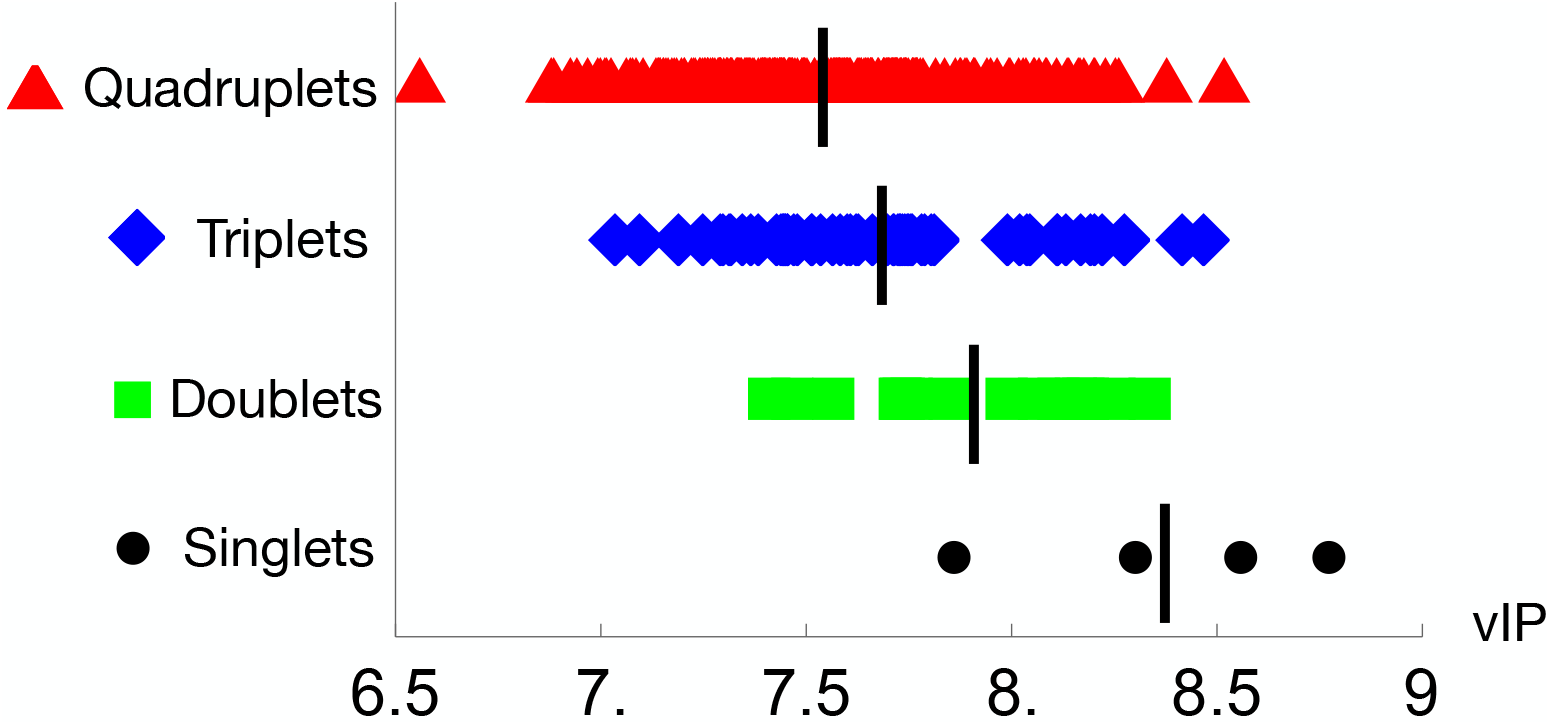
Calculated vIP values at MP2/6-31G* level of theory (in eV) for all singlet, pair, triplet and quadruplet base pair motifs. The mean values are indicated as vertical bars.

**Table 3:** Calculated vIP values of triplets of successive nucleobases in B1 conformation (in eV). The linear correlation coefficient *r* between the calculated vIP and log 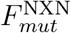, the logarithm of the mutation frequency, is equal to −0.60 (*p*-value 1 × 10^−7^).

| MP2 |  |  |  | 6-31G* |  |  |  |
| --- | --- | --- | --- | --- | --- | --- | --- |
| GGG | 7.09 | AGG | 7.19 | CGG | 7.03 | TGG | 7.34 |
| GGA | 7.44 | AGA | 7.44 | CGA | 7.29 | TGA | 7.60 |
| GGC | 7.45 | AGC | 7.47 | CGC | 7.32 | TGC | 7.63 |
| GGT | 7.43 | AGT | 7.45 | CGT | 7.30 | TGT | 7.61 |
| GAG | 7.58 | AAG | 7.45 | CAG | 7.38 | TAG | 7.51 |
| GAA | 7.71 | AAA | 7.53 | CAA | 7.44 | TAA | 7.66 |
| GAC | 7.73 | AAC | 7.75 | CAC | 7.81 | TAC | 8.11 |
| GAT | 7.69 | AAT | 7.71 | CAT | 7.74 | TAT | 8.04 |
| GCG | 7.48 | ACG | 7.31 | CCG | 7.25 | TCG | 7.36 |
| GCA | 7.76 | ACA | 7.75 | CCA | 7.68 | TCA | 7.80 |
| GCC | 7.78 | ACC | 8.27 | CCC | 7.45 | TCC | 8.02 |
| GCT | 7.73 | ACT | 8.22 | CCT | 7.51 | TCT | 7.73 |
| GTG | 7.79 | ATG | 7.62 | CTG | 7.56 | TTG | 7.68 |
| GTA | 7.72 | ATA | 8.04 | CTA | 7.99 | TTA | 8.11 |
| GTC | 7.75 | ATC | 8.19 | CTC | 8.42 | TTC | 8.17 |
| GTT | 7.70 | ATT | 8.13 | CTT | 8.20 | TTT | 8.47 |

In the same way as we did for pair motifs in the previous section, we calculated the correlation between the vIP values of base triplets and quadruplets with log 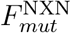 and log 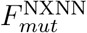, respectively, i.e. the logarithm of the frequencies of all possible base triplet and quadruplet motifs with the middle base mutated, as obtained from the ClinVar database. ^47^ The correlation coefficient decreases somewhat, from −0.80 for pairs, −0.60 for triplets and −0.50 for quadruplets, but the statistical significance increases, as shown by the *p*-values of 2 × 10^−4^, 1 × 10^−7^ and 7 × 10^−18^. These correlations are thus highly statistically significant.

### Methylated Cyt

The epigenetic process of DNA methylation involves adding methyl groups to specific nucleobases and thus modifies the activity of the DNA without altering its sequence. The most common methylation process targets cytosine and modifies it into 5-methylcytosine (5mC or M). Cytosine methylation and demethylation are highly regulated processes, which occur basically across all species where they play key roles in many biological processes. ^48^

Because of its importance, we computed the vIP of 5mC using MP2/6-31G* theory. As seen in Table 4, it is equal to 8.2 eV, and thus lower than the vIP of Cyt (8.6 eV) in agreement with earlier studies^49,50^, and even slightly lower than the vIP of Ade (8.3 eV) (Table 1). The ranking from lowest to highest vIPs is thus: Gua, 5mC, Ade, Cyt, Thy. Gua thus remains the best electron hole trap, but 5mC appears as the second best. Cytosine methylation is therefore expected to impact long-range charge transport through DNA as well as mutability properties.

**Table 4:**
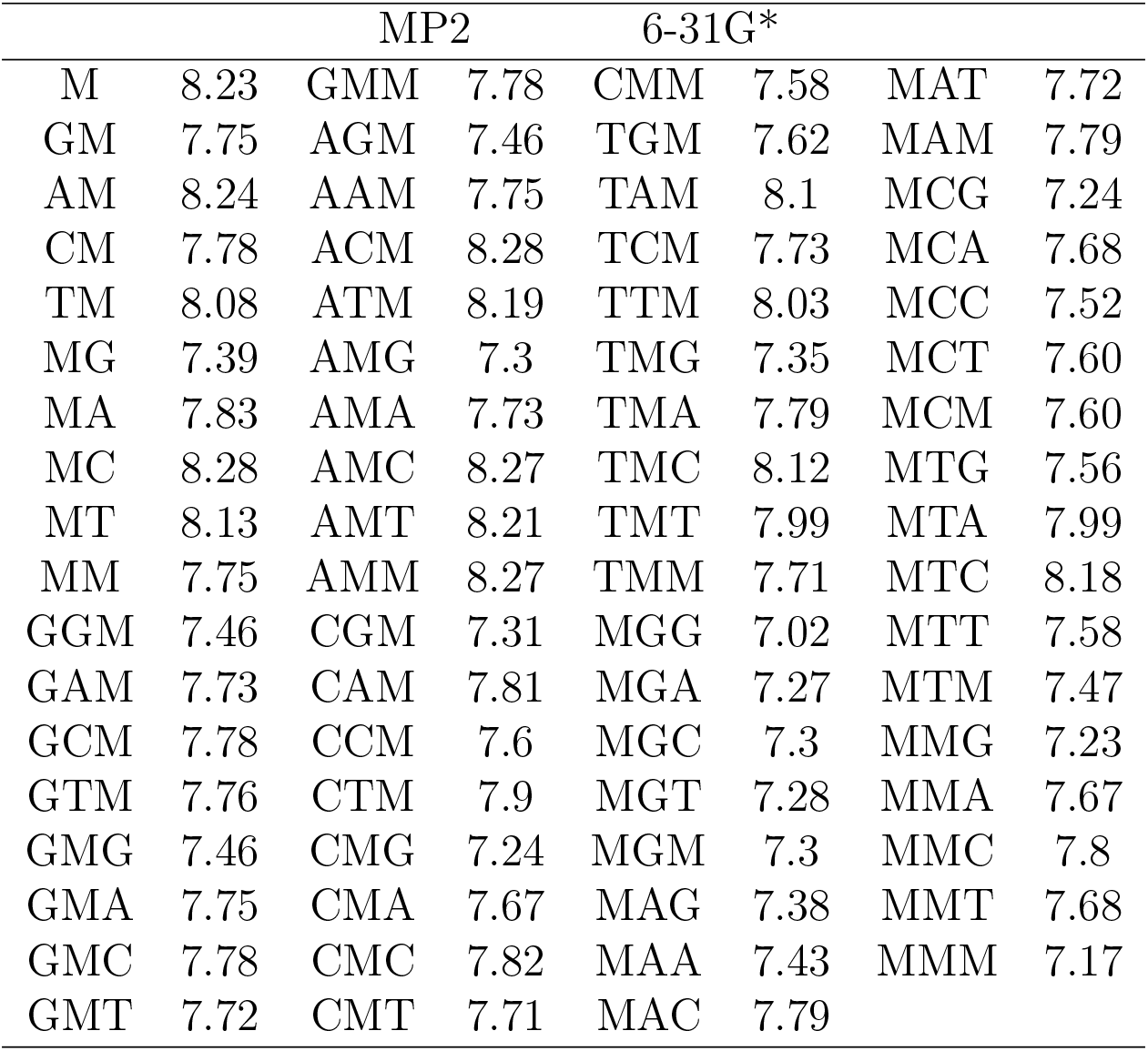
Calculated vIP values (in eV) of 5mC and of pairs and triplets of successive nucleobases in B1 conformation containing at least one 5mC (M).

We also calculated the vIPs of all pairs, triplets and quadruplets containing at least one 5mC (Table 4 and Supplementary Table S5). By comparing these results with Tables 2, 3 and S4, we see that Cyt methylation tends to lower the vIP of the DNA stacks. As lower vIPs are associated with higher mutability, we expect that DNA regions in which cytosines are methylated are more subject to mutation,

To check this, we correlated the vIP values of all pairs, triplets and quadruplets in which we considered all cytosines to be methylated, with the logarithm of the frequency of synonymous variants in ClinVar; note that it is not known whether the cytosines are methylated or not. By doing that, we found better correlation coefficients when assuming all Cyt bases to be methylated rather than unmethylated. As shown in Fig. 3, we found indeed that *r* improves from −0.80 to −0.90 for pairs (*p*-value 2 × 10^−4^ and 2 × 10^−6^), from −0.60 to −0.64 for triplets (*p*-value 1 × 10^−7^ and 1 × 10^−8^), and −0.50 to −0.54 for quadruplets (*p*-value 7 × 10^−18^ and 2 × 10^−20^). This result suggests that many of the cytosines in the genome are methylated, and that the mutability of motifs with methylated cytosines is on the average higher than with unmethylated cytosines, which is indeed observed.^51^ It also supports the hypothesis that the higher mutability of 5mC compared to Cyt is due to its lower vIP value.

**Figure 3:**
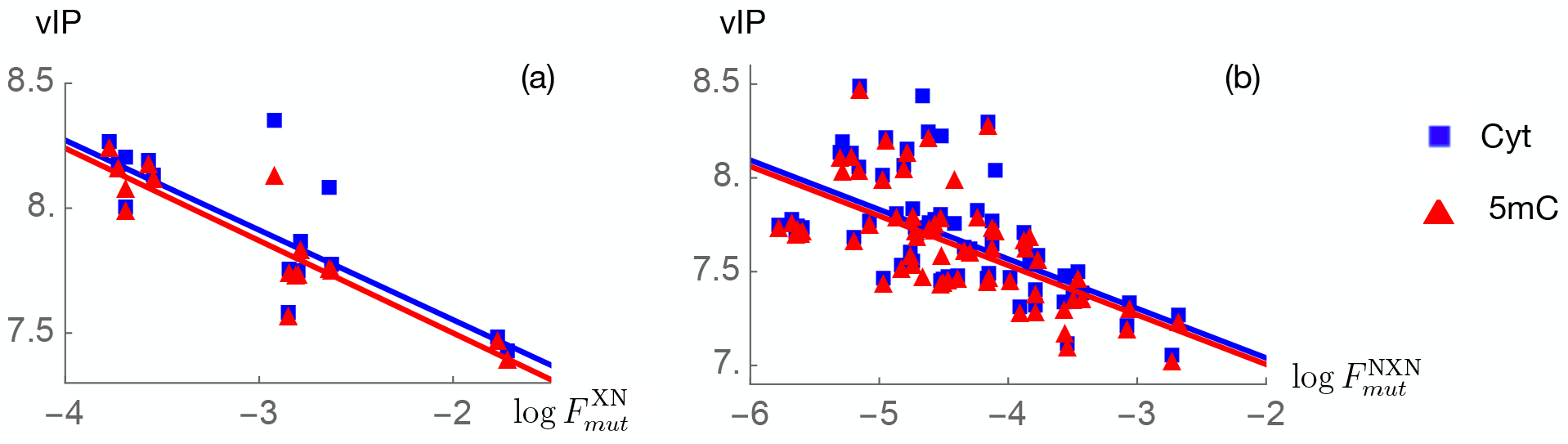
Calculated vIP values at MP2/6-31G* level of theory (in eV) as a function of the logarithm of the mutation frequency, for: (a) all nucleobase pairs of Gua, Ade, Thy and Cyt (blue) and of Gua, Ade, Thy and 5mC (red); (b) all nucleobase triplets of Gua, Ade, Thy and Cyt (blue) and of Gua, Ade, Thy and 5mC (red). The linear regression lines are drawn.

### vIPer for long single-stranded DNA stacks

We constructed a simple mathematical model based on the vIP values of single bases, doublets, triplets and quadruplets, calculated at MP2/6-31G* level of theory, which were combined to predict the vIP of single-stranded nucleobase sequences of any length, containing arbirary combinations of Gua, Ade, Thy, Cyt and 5mC. To test the model, we also calculated at MP2/6-31G* level the vIP values of 249 randomly chosen quintuplets and 42 sextuplets; we call these our test set.

We started by noticing that the vIP values of nucleobase stacks generally decrease as their length *L* increases (see Figs 2 and 4.a). This is expected, as *π*-*π* stacks of increased length allow for more efficient charge delocalization and are easier to ionize. We chose an exponential decay function to model the average stack length dependence of the vIPs:

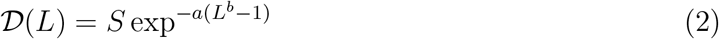

where *S* is the average of the vIP values of the five single bases Gua, Ade, Thy, Cyt and 5mC. The parameters *a* and *b* were identified in order to minimize the RMSE between 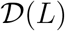 and the average vIP of all nucleobase stack motifs of given length *L* = 1,… 4. The resulting 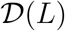 function is plotted in Fig. 4.a. For *L* = 1, we have 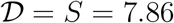; for *L* > 1, 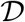 decays exponentially; and for very large *L*, 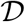 is around 0.6.

**Figure 4:**
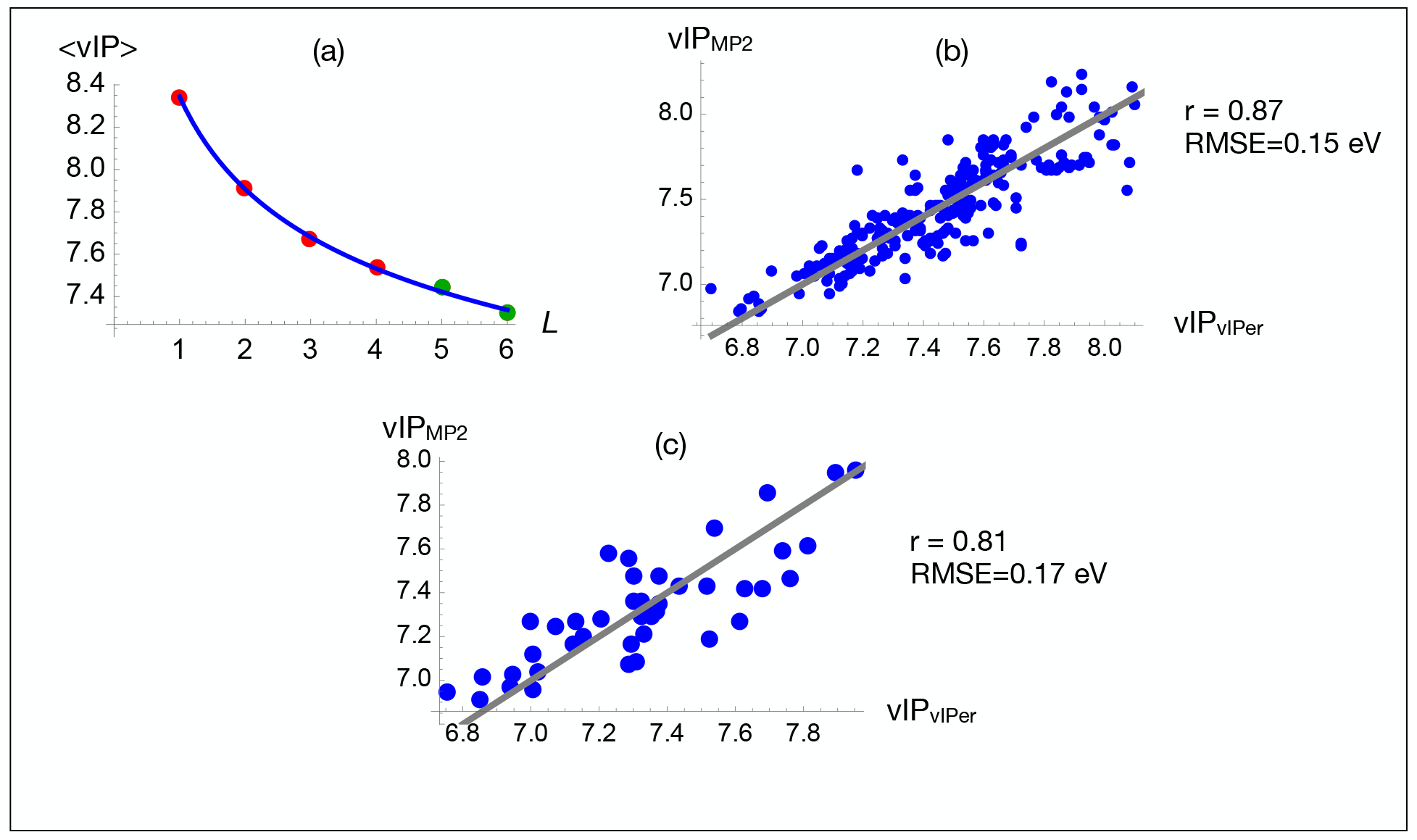
Prediction of vIP values for long single-stranded DNA sequences (a) Exponential decay of the average vIP value of nucleobase sequences of given length *L* as a function of *L*. The points correspond to the vIP values calculated at MP2/6-31G* level; for *L* = 1,… 4 (red points) the average is over all possible nucleobase motifs and and for *L* = 4, 5 (green points), it is over all randomly chosen motifs included in the test set. The blue curve is the exponential decay curve 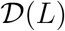 defined in Eq. (2) fitted on the red points *L* = 1,… 4. The green points that correspond to *L* = 5, 6 are not used for fitting. (b)-(c) The vIP values calculated at MP2/6-31G* level as a function of the vIP values calculated by our model defined by Eq. (3) for the test set of quintuplet motifs (b) and sextuplet motifs (c). The Pearson correlation coefficient *r* and RMSE (in eV) are given in the plots. The lines in (b) and (c) are the bisectors of quadrants I and III and correspond to perfect predictions.

The vIP of a given nucleobase sequence (*x*_1_….*x_L_*) of length *L*, where each *x_i_* is one of the five nucleobases Gua, Ade, Thy, Cyt and 5mC, is then expressed as a recursive function of 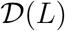 and of the vIP values of the two overlapping sequences of length *L* – 1, i.e. (*x*_1_….*x*_*L*–1_) and (*x*_2_….*x_L_*), as:

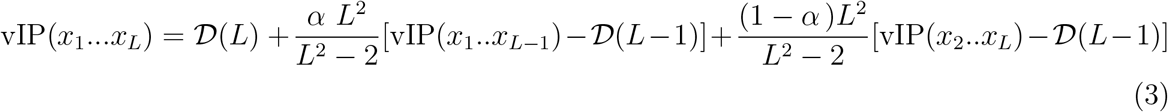

where *L* > 1. The parameter *α* = 0.4 was identified by minimizing the RMSE between the vIP values of all possible quadruplet stacks either calculated at MP2/6-31G* level or estimated via Eq. (3) for *L* = 4. The factor *L*^2^/(*L*^2^ – 2) was introduced to avoid the shrinking of the standard deviation of the distribution of vIP values for long DNA stacks.

Note that we also tried other recursive models, which contained additional free parameters to optimize or were based on more complex model structures. However, we found that the selected model defined in Eq. (3), which has a very simple structure and only α as free parameter, in which the vIP of an *L*-tuplet is basically derived from the vIP of the two overlapping (*L*-1)-tuplets, is less prone to biases and reaches the best score.

An important observation from our model is that the value of the α parameter is lower than 0.5, which means that the vIP value of the (*L*-1)-tuplet near the 3’ end contributes more than the vIP of the (*L*-1)-tuplet near the 5’ end to the vIP value of the *L*-tuplet. This asymmetry can be related to the 5’→3’ directional preference of the hole transport through the DNA stack.

To check the performance of the model, we applied it to calculate the vIP values of the quintuplet and sextuplet motifs of our test set. As shown in Fig. 4.b-c, the results are very good with Pearson correlation coefficients between our model predictions and our quantum chemistry calculations of 0.87 (*p*-value 9 × 10^−79^) and 0.81 (*p*-value 2 × 10^−11^) for the quintuplets and sextuplets, respectively, and a RMSE of 0.15 and 0.17 eV.

To make the results of our prediction model available to the scientific community, we developed a python package called vIPer, which can be freely dowloaded from our GitHub repository (github.com/3BioCompBio/vIPer). For nucleobase sequences of length between one and four, vIPer outputs the vIP value calculated at MP2/6-31G*. For sequences longer than four bases, vIPer uses the recursive model of Eq. (3). vIPer is simple to use: the user submits a nucleobase sequence and gets the vIP values predicted by the model for the input sequence as well as the average over the two complementary strands. It has the advantage of avoiding intense quantum chemistry calculations to estimate the vIP of long nucleobase stacks, while maintaining very good precision. Examples of predicted vIP values for DNA sequences of interest are shown in Table 5.

**Table 5:** vIP values predicted by vIPer (in eV) of examples of long single-base DNA stacks in B1 conformation: telomers,^52^ pericentromeric satellite repeats^53^ and methylated or unmethylated CpG islands. ^54^ The vIP column contains the predicted vIP of the motif and the <vIP> column, the predicted vIP averaged over the two complementary strands.

| Type | organisms | DNA motif | vIP | <vIP> |
| --- | --- | --- | --- | --- |
| Telomers | vertebrates | (TTAGGG)<sub>100</sub> | 5.50 | 5.82 |
| Telomers | plants | (TTTAGGG)<sub>100</sub> | 5.67 | 5.91 |
| pericentromeric satellite | *Drosophila* | (AATAT)<sub>35</sub> | 6.55 | 6.73 |
| pericentromeric satellite | *Drosophila* | (AAGAGAG)<sub>25</sub> | 6.20 | 6.64 |
| pericentromeric satellite | *Drosophila* | (AATAACATAG)<sub>8</sub> | 6.48 | 6.69 |
| CpG island | vertebrates | (CG)<sub>500</sub> | 5.59 | 5.59 |
| methylated CpG | vertebrates | (MG)<sub>500</sub> | 5.56 | 5.56 |

### Validation of vIPer on experimental data

To further validate the vIPer model, we compared its predictions with experimental data. We first considered photoinduced DNA cleavage experiments. ^20^ In these experiments, one-electron oxidation of G- and GG-containing DNA segments were examined as a function of the flanking sequence through the measurement of their relative reactivity (*k_rel_*). The nucleobase motif that is varied is the 5-tuplet TNNNT, where N denotes any nucleobase; the full 30-tuplet sequence is specified in Table 6. We compared *k^rel^* with vIPer’s vIP values of the 5-tuplet and 30-tuplet sequences and found excellent Pearson correlation coefficients of −0.91 and −0.93, respectively; these values are statistically significant (*p*-values 0.00004 and 9 × 10^−6^).

**Table 6:** Comparison between the measured relative reactivity *k_rel_* in photoinduced DNA cleavage experiments of 30-tuplet sequences^20^ and their vIP value (in eV) calculated using vIPer. The DNA sequences are CGTACTCTTTGGTGGG**TNNNT**TCTTTCTAT, where N denotes any base. The tested TNNNT quintuplet motifs are given in column 1. The Pearson correlation coefficient *r* is between *k_rel_* and vIP values.

| Motifs | $k_{rel}^{20}$ | vIP | |
| --- | --- | --- | --- |
|  |  | 5-tuplet | 30-tuplet |
| TGGGT | 2.7 | 6.692 | 6.120 |
| TAGGT | 1.0 | 7.004 | 6.346 |
| TTGGT | 1.0 | 7.174 | 6.464 |
| TCGGT | 2.0 | 6.840 | 6.236 |
| TGGAT | 0.8 | 7.257 | 6.496 |
| TAGAT | 0.4 | 7.283 | 6.566 |
| TTGAT | 0.3 | 7.455 | 6.684 |
| TCGAT | 0.4 | 7.121 | 6.457 |
| TGGTT | 0.9 | 7.243 | 6.531 |
| TAGTT | 0.2 | 7.289 | 6.609 |
| TGGCT | 0.7 | 7.276 | 6.523 |
| TAGCT | 0.1 | 7.316 | 6.599 |
| $r$ | | -0.91 | -0.93 |
| $p$-value | | 0.00004 | $9 \times 10^{-6}$ |

As an additional validation test, we compared vIPer’s predictions with experiments in which the effects of the DNA sequence on charge transport were analyzed using cyclic voltammetry of daunomycin cross-linked with different palindromic DNA duplexes. ^55^ Based on the empirical linear relationship between the ionization potential and the peak potential for oxidation *V_ox_*,^56,57^ we compared the latter with the calculated vIP values for the considered duplexes, as shown in Table 7. The Pearson correlation coefficient between *V_ox_* and vIP values is equal to 0.89 and statistically significant (*p*-value 0.003). This result further demonstrates the quality of vIPer.

**Table 7:**
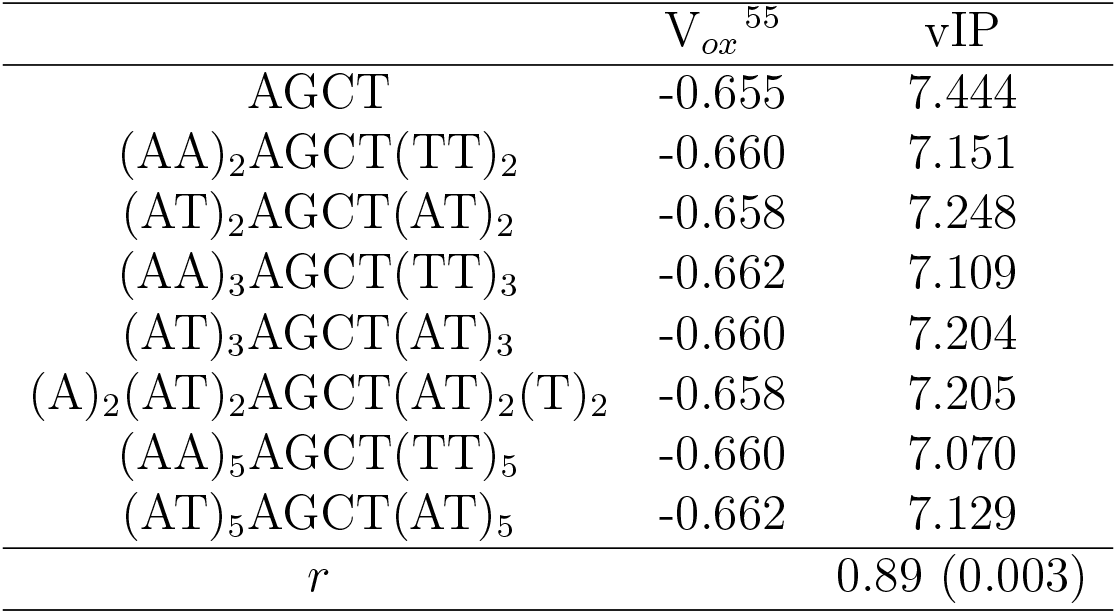
Comparison between the vIP (in eV) calculated by vIPer and the peak potential for oxidation *V_ox_* (in V) measured by cyclic voltammetry on the palindromic DNA duplexes of column 1.^55^ The Pearson correlation coefficient *r* is between vIP and *V_ox_*; the *p*-value is in parentheses.

| | $V_{ox}$<sup>55</sup> | vIP |
| --- | --- | --- |
| AGCT | -0.655 | 7.444 |
| (AA)<sub>2</sub>AGCT(TT)<sub>2</sub> | -0.660 | 7.151 |
| (AT)<sub>2</sub>AGCT(AT)<sub>2</sub> | -0.658 | 7.248 |
| (AA)<sub>3</sub>AGCT(TT)<sub>3</sub> | -0.662 | 7.109 |
| (AT)<sub>3</sub>AGCT(AT)<sub>3</sub> | -0.660 | 7.204 |
| (A)<sub>2</sub>(AT)<sub>2</sub>AGCT(AT)<sub>2</sub>(T)<sub>2</sub> | -0.658 | 7.205 |
| (AA)<sub>5</sub>AGCT(TT)<sub>5</sub> | -0.660 | 7.070 |
| (AT)<sub>5</sub>AGCT(AT)<sub>5</sub> | -0.662 | 7.129 |
| $r$ | | 0.89 (0.003) |

## Conclusion

We performed in this paper a huge number of quantum chemistry calculations at different levels of theory and using various basis sets in view of estimating the vIP values of all possible DNA stacks of arbitrary lengths. This is as such an important achievement. Indeed, vIPer is the very first automatic vIP calculator of single stranded DNA molecules of any sequence and any length, and moreover, it is very fast as it computes the vIP values in just a few seconds. We would like to underline that we found very good correlations between vIPer predictions and experimental cyclic voltammetry and photoinduced one-electron oxidation measurements.

Another point we would like to emphasize is the importance of considering the *π*-orbital overlap between stacked nucleobases and the associated dispersion energy contributions when performing vIP calculations. Indeed, we compared the MP2/6-31G* vIP values of all pairs, triplets and quadruplets obtained by calculating them either from the complete nucleobase stacks or as the average of the vIP of their constituting nucleobases. The former method is clearly superior. Indeed, we found that the Pearson correlation coefficients between the vIP and the logarithm of the mutations frequency (log *F_mut_*) deteriorate when passing from the former to the latter method from −0.80 to −0.70 for pairs, from −0.61 to −0.32 for triplets and from −0.50 to −0.23 for quadruplets.

vIPer has a whole series of applications. Indeed, the estimation of the electronic properties of DNA can be crucial in biomedical investigations aiming, for example, to predict and control the mutability of specific genome regions or other vIP-dependent biological processes. It is also important in biotechnological applications in which DNA molecules are used for their unique properties such as the construction of molecular wires or data storage systems made from DNA. ^58^

We would like to stress that understanding the charge transport properties of DNA is still elusive and that even experimental studies have so far not provided definitive answers. Computational analyses are therefore of primary importance in this context for suggesting new hypotheses and guiding new research.

Even though vIPer is certainly a first important step towards accurate vIP estimations of DNA molecules, it is based on a number of approximations that are important to discuss:

- The use of gold-standard quantum chemistry methods such as CCSD(T)^46^ rather than MP2 would be a clear improvement, but they are currently too time- and memory-consuming to be applied to all possible pair to quadruplet motifs.
- We omitted the sugar-phosphate backbone in all our calculations, as their inclusion would require too much additional computer time and memory. Another justification of this choice is that earlier calculations on Cyt and Thy motifs indicated that the vIP is strongly affected by the presence of the sugar and phosphate moieties in gas phase, but much less upon bulk hydration, due to the screening by the aqueous solvent.^59^ Moreover, quantum chemistry calculations and experimental data showed that the lowest ionization pathway comes from the nucleobases rather than the sugar-phosphate backbone.^59,60^
- DNA environment makes the vIP estimation very challenging. Here we disregarded the solvent and performed all calculations in gas phase, although solvation is known as impacting the electronic properties of DNA molecules. ^61^ This is of course an approximation, but appears to be justified when comparing vIP values of various nucleobase stack sequences. Indeed, calculations on individual nucleobases highlighted the effect of the solvent in lowering the vIP values while maintaining the relative ordering between the bases.^62,63^ Moreover, as discussed above, dropping both the sugar-phosphate backbone and the solvent have opposite effects that tend to cancel out. ^59^

Finally note that the vIP values depend significantly on the type of DNA conformation. Indeed, we clearly showed here that the vIPs differ according to the type of B-conformation we considered (B1 and B2), and it was previously shown that it is even more so when considering A- or Z-conformations instead of B-conformations.^23^ We will in the future extend the vIPer algorithm to make it able to predict the vIP of other DNA conformations and introduce additional parameters in the model that take into account the high flexibility of DNA.

## Acknowledgments

We thank Emilie Cauët and Nicolas Callebaut for interesting discussions at the beginning of the project. We acknowledge financial support from the FNRS - Fund for Scientific Research through a PDR Research Project; MR is FNRS Research Director.

## Data and Software Availability

All quantum chemistry calculations were performed using the Gaussian 16 suite.^40^ The geometry of the single nucleobases and nucleobase stacks were constructed using the software package x3DNA-DSSR,^41^ and the nucleobases with optimized geometry were superimposed onto the DNA stacks using the U3BEST algorithm.^44^ The python code of the vIPer model that estimates the vIP of single-stranded DNA molecules of any length in B conformation is freely available on our github repository github.com/3BioCompBio/vIPer.

