## Supplemental Data 1 for "Estimating the vertical ionization potential of single-stranded DNA molecules"

**Table S1.** Calculated and experimental vIP values of single nucleobases in eV.
**Table S2.** Calculated vIP values of pairs of successive nucleobases in B1 conformation (in eV).
**Table S3.** Calculated vIP values of pairs of successive nucleobases in B2 conformation (eV).
**Table S4.** Calculated vIP values of all quadruplets of successive nucleobases in B1 conformation (in eV) containing Gua, Ade, Cyt or Thy.
**Table S5.** Calculated vIP values of quadruplets of successive nucleobases in B1 conformation (in eV) containing at least one 5mC (M).

| Level of theory | MP2 | | | | Experi |
|---|---|---|---|---|---|
| Basis set | 6-31G* | 6-31G** | 6-31G$^{(.2)}$ | 6-31G$^{(.2)*}$ | mental [1] |
| Gua | 7.86 | 7.89 | 7.99 | 8.04 | 8.24 |
| Ade | 8.30 | 8.34 | 8.37 | 8.44 | 8.44 |
| Cyt | 8.56 | 8.61 | 8.56 | 8.61 | 8.94 |
| Thy | 8.77 | 8.83 | 8.88 | 8.94 | 9.14 |
| RMSE | 0.33 | 0.29 | 0.26 | 0.22 | |
| $r$ | 0.96 | 0.96 | 0.96 | 0.95 | |
| $p$-value | 0.04 | 0.04 | 0.04 | 0.05 | |
| Level of theory | B2-PLYP | | | | Experi |
| Basis set | 6-31G* | 6-31G** | 6-31G$^{(.2)}$ | 6-31G$^{(.2)*}$ | mental |
| Gua | 7.71 | 7.71 | 7.83 | 7.82 | 8.24 |
| Ade | 8.04 | 8.04 | 8.17 | 8.15 | 8.44 |
| Cyt | 8.43 | 8.44 | 8.54 | 8.54 | 8.94 |
| Thy | 8.72 | 8.73 | 8.83 | 8.84 | 9.14 |
| RMSE | 0.47 | 0.46 | 0.35 | 0.36 | |
| $r$ | 0.99 | 0.99 | 0.99 | 0.99 | |
| $p$-value | 0.009 | 0.009 | 0.01 | 0.01 | |
| Level of theory | mPW2-PLYP | | | | Experi |
| Basis set | 6-31G* | 6-31G** | 6-31G$^{(.2)}$ | 6-31G$^{(.2)*}$ | mental |
| Gua | 7.75 | 7.75 | 7.86 | 7.85 | 8.24 |
| Ade | 8.08 | 8.07 | 8.20 | 8.18 | 8.44 |
| Cyt | 8.49 | 8.49 | 8.59 | 8.58 | 8.94 |
| Thy | 8.77 | 8.78 | 8.88 | 8.88 | 9.14 |
| RMSE | 0.42 | 0.42 | 0.31 | 0.32 | |
| $r$ | 0.99 | 0.99 | 0.99 | 0.99 | |
| $p$-value | 0.008 | 0.008 | 0.01 | 0.008 | |
| Level of theory | PBE0-DH | | | | Experi |
| Basis set | 6-31G* | 6-31G** | 6-31G$^{(.2)}$ | 6-31G$^{(.2)*}$ | mental |
| Gua | 7.94 | 7.93 | 8.00 | 7.97 | 8.24 |
| Ade | 8.27 | 8.25 | 8.35 | 8.31 | 8.44 |
| Cyt | 8.73 | 8.72 | 8.79 | 8.77 | 8.94 |
| Thy | 8.98 | 8.98 | 9.02 | 9.01 | 9.14 |
| RMSE | 0.22 | 0.23 | 0.16 | 0.18 | |
| $r$ | 1.00 | 1.00 | 0.99 | 0.99 | |
| $p$-value | 0.005 | 0.004 | 0.007 | 0.005 | |

Table S1. Calculated and experimental vIP values of single nucleobases in eV. The root mean square error (RMSE) and the Pearson correlation coefficient $r$ (with associated p-values) are computed between the calculated and experimental [1] vIP values.

| Level of theory | MP2 | | | | B2-PLYP | | | |
|---|---|---|---|---|---|---|---|---|
| Basis set | 6-31G* | 6-31G** | 6-31G$^{(.2)}$ | 6-31G$^{(.2)*}$ | 6-31G* | 6-31G** | 6-31G$^{(.2)}$ | 6-31G$^{(.2)*}$ |
| GG | 7.47 | 7.50 | 7.61 | 7.66 | 7.27 | 7.27 | 7.42 | 7.41 |
| GA | 7.73 | 7.75 | 7.78 | 7.85 | 7.51 | 7.50 | 7.64 | 7.62 |
| GC | 7.76 | 7.78 | 7.81 | 7.87 | 7.58 | 7.57 | 7.69 | 7.67 |
| GT | 7.74 | 7.76 | 7.86 | 7.91 | 7.57 | 7.57 | 7.69 | 7.68 |
| AG | 7.57 | 7.61 | 7.71 | 7.77 | 7.26 | 7.27 | 7.48 | 7.44 |
| AA | 7.99 | 8.04 | 7.77 | 7.86 | 7.50 | 7.51 | 7.67 | 7.64 |
| AC | 8.25 | 8.29 | 8.25 | 8.34 | 7.96 | 7.95 | 8.08 | 8.05 |
| AT | 8.17 | 8.22 | 8.24 | 8.32 | 7.86 | 7.87 | 8.01 | 8.00 |
| CG | 7.41 | 7.45 | 7.55 | 7.61 | 7.23 | 7.24 | 7.38 | 7.37 |
| CA | 7.85 | 7.90 | 7.94 | 8.02 | 7.56 | 7.57 | 7.70 | 7.69 |
| CC | 8.07 | 8.12 | 8.04 | 8.09 | 7.88 | 7.88 | 8.00 | 8.00 |
| CT | 8.34 | 7.57 | 8.42 | 8.48 | 7.89 | 7.90 | 8.02 | 8.03 |
| TG | 7.73 | 7.76 | 7.85 | 7.90 | 7.52 | 7.53 | 7.67 | 7.66 |
| TA | 8.16 | 8.20 | 8.21 | 8.29 | 7.85 | 7.86 | 7.98 | 7.97 |
| TC | 8.19 | 8.24 | 8.16 | 8.23 | 8.14 | 8.15 | 8.27 | 8.27 |
| TT | 8.11 | 8.18 | 8.18 | 8.24 | 8.13 | 8.15 | 8.25 | 8.26 |
| $r$ | -0.80 | -0.85 | -0.70 | -0.72 | -0.73 | -0.73 | -0.75 | -0.73 |
| $p$-value | $2 \times 10^{-4}$ | $3 \times 10^{-5}$ | $2 \times 10^{-3}$ | $2 \times 10^{-3}$ | $1 \times 10^{-3}$ | $1 \times 10^{-3}$ | $8 \times 10^{-4}$ | $1 \times 10^{-3}$ |

| Level of theory | mPW2-PLYP | | | | PBE0-DH | | | |
|---|---|---|---|---|---|---|---|---|
| Basis set | 6-31G* | 6-31G** | 6-31G$^{(.2)}$ | 6-31G$^{(.2)*}$ | 6-31G* | 6-31G** | 6-31G$^{(.2)}$ | 6-31G$^{(.2)*}$ |
| GG | 7.32 | 7.32 | 7.45 | 7.43 | 7.47 | 7.45 | 7.56 | 7.53 |
| GA | 7.56 | 7.54 | 7.67 | 7.65 | 7.69 | 7.66 | 7.80 | 7.71 |
| GC | 7.61 | 7.60 | 7.72 | 7.69 | 7.78 | 7.74 | 7.86 | 7.81 |
| GT | 7.61 | 7.61 | 7.72 | 7.71 | 7.79 | 7.78 | 7.85 | 7.82 |
| AG | 7.34 | 7.45 | 7.56 | 7.54 | 7.52 | 7.51 | 7.62 | 7.61 |
| AA | 7.58 | 7.57 | 7.74 | 7.71 | 7.78 | 7.76 | 7.89 | 7.84 |
| AC | 8.00 | 7.98 | 8.12 | 8.08 | 8.17 | 8.13 | 8.26 | 8.19 |
| AT | 7.92 | 7.92 | 8.06 | 8.04 | 8.09 | 8.08 | 8.18 | 8.14 |
| CG | 7.28 | 7.28 | 7.40 | 7.39 | 7.44 | 7.44 | 7.53 | 7.50 |
| CA | 7.60 | 7.61 | 7.73 | 7.71 | 7.78 | 7.78 | 7.87 | 7.83 |
| CC | 7.99 | 7.99 | 8.10 | 8.10 | 8.15 | 8.14 | 8.24 | 8.21 |
| CT | 7.98 | 7.99 | 8.11 | 8.11 | 8.24 | 8.25 | 8.32 | 8.31 |
| TG | 7.58 | 7.58 | 7.70 | 7.69 | 7.75 | 7.73 | 7.83 | 7.80 |
| TA | 7.90 | 7.91 | 8.02 | 8.00 | 8.08 | 8.06 | 8.16 | 8.12 |
| TC | 8.24 | 8.24 | 8.36 | 8.36 | 8.46 | 8.45 | 8.54 | 8.51 |
| TT | 8.22 | 8.23 | 8.34 | 8.34 | 8.48 | 8.49 | 8.54 | 8.53 |
| $r$ | -0.72 | -0.73 | -0.74 | -0.73 | -0.71 | -0.70 | -0.72 | -0.71 |
| $p$-value | $2 \times 10^{-3}$ | $1 \times 10^{-3}$ | $9 \times 10^{-4}$ | $1 \times 10^{-3}$ | $2 \times 10^{-3}$ | $2 \times 10^{-3}$ | $2 \times 10^{-3}$ | $2 \times 10^{-3}$ |

Table S2. Calculated vIP values of pairs of successive nucleobases in B1 conformation (in eV). The Pearson correlation coefficient $r$ and associated $p$-values are computed between the calculated vIP and $\log F_{mut}^{\mathrm{XN}}$, the logarithm of the mutation frequency observed in the ClinVar database [2].

| Level of theory | MP2 | | | |
|---|---|---|---|---|
| Basis set | 6-31G* | 6-31G** | 6-31G$^{(.2)}$ | 6-31G$^{(.2)*}$ |
| GG | 7.48 | 7.51 | 7.63 | 7.68 |
| GA | 7.73 | 7.75 | 7.78 | 7.85 |
| GC | 7.74 | 7.76 | 7.80 | 7.86 |
| GT | 7.72 | 7.75 | 7.85 | 7.90 |
| AG | 7.58 | 7.63 | 7.73 | 7.78 |
| AA | 7.73 | 7.77 | 7.72 | 7.81 |
| AC | 8.23 | 8.26 | 8.23 | 8.31 |
| AT | 8.17 | 8.21 | 8.23 | 8.31 |
| CG | 7.46 | 7.50 | 7.61 | 7.67 |
| CA | 7.89 | 7.95 | 7.99 | 8.06 |
| CC | 8.15 | 8.21 | 8.11 | 8.17 |
| CT | 8.43 | 8.50 | 8.51 | 8.58 |
| TG | 7.76 | 7.80 | 7.89 | 7.94 |
| TA | 8.18 | 8.23 | 8.24 | 8.31 |
| TC | 8.83 | 8.89 | 8.84 | 8.91 |
| TT | 8.17 | 8.24 | 8.24 | 8.30 |
| $r$ | -0.67 | -0.66 | -0.63 | -0.64 |
| $p$-value | $5 \times 10^{-3}$ | $5 \times 10^{-3}$ | $9 \times 10^{-3}$ | $7 \times 10^{-3}$ |

Table S3. Calculated vIP values of pairs of successive nucleobases in B2 conformation (eV). The Pearson correlation coefficient $r$ and associated $p$-values are computed between the calculated vIP and $\log F_{mut}^{\mathrm{XN}}$, the logarithm of the mutation frequency observed in the ClinVar database [2].

| | | | | | MP2 | | | 6-31G* | | | | | | | |
|---|---|---|---|---|---|---|---|---|---|---|---|---|---|---|---|
| GGGG | 6.97 | GCGG | 7.10 | AGGG | 6.93 | ACGG | 6.94 | CGGG | 6.89 | CCGG | 6.88 | TGGG | 6.56 | TCGG | 6.99 |
| GGGA | 7.07 | GCGA | 7.36 | AGGA | 7.16 | ACGA | 7.20 | CGGA | 7.01 | CCGA | 7.13 | TGGA | 7.32 | TCGA | 7.25 |
| GGGC | 7.09 | GCGC | 7.38 | AGGC | 7.18 | ACGC | 7.22 | CGGC | 7.02 | CCGC | 7.16 | TGGC | 7.33 | TCGC | 7.27 |
| GGGT | 7.06 | GCGT | 7.36 | AGGT | 7.15 | ACGT | 7.20 | CGGT | 7.00 | CCGT | 7.14 | TGGT | 7.31 | TCGT | 7.25 |
| GGAG | 7.37 | GCAG | 7.42 | AGAG | 7.57 | ACAG | 7.34 | CGAG | 7.15 | CCAG | 7.31 | TGAG | 7.46 | TCAG | 7.36 |
| GGAA | 7.44 | GCAA | 7.76 | AGAA | 7.43 | ACAA | 7.37 | CGAA | 7.28 | CCAA | 7.33 | TGAA | 7.58 | TCAA | 7.41 |
| GGAC | 7.44 | GCAC | 7.76 | AGAC | 7.44 | ACAC | 7.71 | CGAC | 7.29 | CCAC | 7.65 | TGAC | 7.60 | TCAC | 7.77 |
| GGAT | 7.42 | GCAT | 7.74 | AGAT | 7.41 | ACAT | 7.64 | CGAT | 7.26 | CCAT | 7.58 | TGAT | 7.57 | TCAT | 7.70 |
| GGCG | 7.39 | GCCG | 7.29 | AGCG | 7.43 | ACCG | 7.20 | CGCG | 7.40 | CCCG | 7.17 | TGCG | 7.46 | TCCG | 7.23 |
| GGCA | 7.46 | GCCA | 7.72 | AGCA | 7.47 | ACCA | 7.63 | CGCA | 7.32 | CCCA | 7.60 | TGCA | 7.63 | TCCA | 7.66 |
| GGCC | 7.47 | GCCC | 7.74 | AGCC | 7.49 | ACCC | 7.64 | CGCC | 7.34 | CCCC | 7.31 | TGCC | 7.65 | TCCC | 7.50 |
| GGCT | 7.44 | GCCT | 7.77 | AGCT | 7.44 | ACCT | 8.26 | CGCT | 7.29 | CCCT | 7.39 | TGCT | 7.60 | TCCT | 7.48 |
| GGTG | 7.35 | GCTG | 7.60 | AGTG | 7.14 | ACTG | 7.52 | CGTG | 7.14 | CCTG | 7.49 | TGTG | 7.77 | TCTG | 7.54 |
| GGTA | 7.42 | GCTA | 7.73 | AGTA | 7.44 | ACTA | 7.94 | CGTA | 7.29 | CCTA | 7.91 | TGTA | 7.59 | TCTA | 7.96 |
| GGTC | 7.44 | GCTC | 7.74 | AGTC | 7.47 | ACTC | 8.23 | CGTC | 7.32 | CCTC | 7.58 | TGTC | 7.62 | TCTC | 8.38 |
| GGTT | 7.41 | GCTT | 7.71 | AGTT | 7.41 | ACTT | 8.20 | CGTT | 7.26 | CCTT | 8.03 | TGTT | 7.57 | TCTT | 8.17 |
| GAGG | 7.25 | GTGG | 7.40 | AAGG | 7.08 | ATGG | 7.24 | CAGG | 7.01 | CTGG | 7.18 | TAGG | 7.14 | TTGG | 7.30 |
| GAGA | 7.58 | GTGA | 7.66 | AAGA | 7.33 | ATGA | 7.49 | CAGA | 7.27 | CTGA | 7.44 | TAGA | 7.39 | TTGA | 7.55 |
| GAGC | 7.58 | GTGC | 7.58 | AAGC | 7.36 | ATGC | 7.52 | CAGC | 7.29 | CTGC | 7.46 | TAGC | 7.42 | TTGC | 7.58 |
| GAGT | 7.57 | GTGT | 7.56 | AAGT | 7.34 | ATGT | 7.50 | CAGT | 7.27 | CTGT | 7.45 | TAGT | 7.40 | TTGT | 7.56 |
| GAAG | 7.65 | GTAG | 7.55 | AAAG | 7.39 | ATAG | 7.46 | CAAG | 7.36 | CTAG | 7.43 | TAAG | 7.42 | TTAG | 7.49 |
| GAAA | 7.71 | GTAA | 7.72 | AAAA | 7.45 | ATAA | 7.57 | CAAA | 7.24 | CTAA | 7.53 | TAAA | 7.50 | TTAA | 7.91 |
| GAAC | 7.72 | GTAC | 7.73 | AAAC | 7.91 | ATAC | 8.16 | CAAC | 7.77 | CTAC | 7.95 | TAAC | 7.63 | TTAC | 8.07 |
| GAAT | 7.70 | GTAT | 7.71 | AAAT | 7.87 | ATAT | 7.59 | CAAT | 7.74 | CTAT | 7.88 | TAAT | 7.59 | TTAT | 7.99 |
| GACG | 7.35 | GTCG | 7.40 | AACG | 7.26 | ATCG | 7.32 | CACG | 7.23 | CTCG | 7.29 | TACG | 7.29 | TTCG | 7.34 |
| GACA | 7.73 | GTCA | 7.76 | AACA | 7.69 | ATCA | 7.75 | CACA | 7.66 | CTCA | 7.72 | TACA | 7.72 | TTCA | 7.78 |
| GACC | 7.74 | GTCC | 7.77 | AACC | 7.76 | ATCC | 8.20 | CACC | 7.84 | CTCC | 7.69 | TACC | 8.14 | TTCC | 7.89 |
| GACT | 7.71 | GTCT | 7.74 | AACT | 7.73 | ATCT | 8.18 | CACT | 7.78 | CTCT | 7.59 | TACT | 8.08 | TTCT | 7.81 |
| GATG | 7.66 | GTTG | 7.72 | AATG | 7.57 | ATTG | 7.63 | CATG | 7.54 | CTTG | 7.61 | TATG | 7.60 | TTTG | 7.66 |
| GATA | 7.69 | GTTA | 7.69 | AATA | 7.86 | ATTA | 8.06 | CATA | 7.40 | CTTA | 8.03 | TATA | 8.02 | TTTA | 8.09 |
| GATC | 7.70 | GTTC | 7.71 | AATC | 7.72 | ATTC | 8.14 | CATC | 7.76 | CTTC | 8.25 | TATC | 8.05 | TTTC | 8.52 |
| GATT | 7.67 | GTTT | 7.68 | AATT | 7.69 | ATTT | 8.11 | CATT | 7.70 | CTTT | 7.70 | TATT | 8.00 | TTTT | 7.89 |

Table S4. Calculated vIP values of all quadruplets of successive nucleobases in B1 conformation (in eV) containing Gua, Ade, Cyt or Thy. The Pearson correlation coefficient $r$ between the calculated vIP and $\log F_{mut}^{\mathrm{NXNN}}$, the logarithm of the mutation frequency, is equal to -0.50 (p-value $7 \times 10^{-18}$).

| | | | | MP2 | | 6-31G* | | | | | |
|---|---|---|---|---|---|---|---|---|---|---|---|
| GAMT | 7.71 | AAMT | 7.73 | CAMT | 7.78 | TAMT | 8.07 | MGTA | 7.27 | MTGC | 7.46 |
| GAMM | 7.75 | AAMM | 7.76 | CAMM | 7.84 | TAMM | 8.14 | MGTC | 7.30 | MTGT | 7.44 |
| GCGM | 7.38 | ACGM | 7.22 | CCGM | 7.16 | TCGM | 7.27 | MGTT | 7.24 | MTGM | 7.45 |
| GCAM | 7.77 | ACAM | 7.71 | CCAM | 7.65 | TCAM | 7.76 | MGTM | 7.30 | MTAG | 7.43 |
| GCCM | 7.28 | ACCM | 7.18 | CCCM | 7.51 | TCCM | 7.58 | MGMG | 7.38 | MTAA | 7.53 |
| GCTM | 7.74 | ACTM | 8.23 | CCTM | 7.81 | TCTM | 7.87 | MGMA | 7.30 | MTAC | 7.94 |
| GCMG | 7.28 | ACMG | 7.19 | CCMG | 7.16 | TCMG | 7.21 | MGMC | 7.33 | MTAT | 7.87 |
| GCMA | 7.71 | ACMA | 7.62 | CCMA | 7.59 | TCMA | 7.65 | MGMT | 7.27 | MTAM | 7.94 |
| GCMC | 7.73 | ACMC | 7.63 | CCMC | 7.65 | TCMC | 7.78 | MGMM | 7.33 | MTCG | 7.29 |
| GCMT | 7.77 | ACMT | 8.26 | CCMT | 7.54 | TCMT | 7.67 | MAGG | 7.01 | MTCA | 7.72 |
| GCMM | 7.26 | ACMM | 7.17 | CCMM | 7.04 | TCMM | 7.15 | MAGA | 7.26 | MTCC | 8.2 |
| GTGM | 7.58 | ATGM | 7.51 | CTGM | 7.46 | TTGM | 7.57 | MAGC | 7.29 | MTCT | 8.17 |
| GTAM | 7.73 | ATAM | 7.62 | CTAM | 7.94 | TTAM | 8.06 | MAGT | 7.27 | MTCM | 7.64 |
| GTCM | 7.77 | ATCM | 8.20 | CTCM | 7.65 | TTCM | 7.71 | MAGM | 7.28 | MTTG | 7.61 |
| GTTM | 7.71 | ATTM | 8.14 | CTTM | 7.94 | TTTM | 8.00 | MAAG | 7.36 | MTTA | 8.03 |
| GTMG | 7.38 | ATMG | 7.30 | CTMG | 7.27 | TTMG | 7.33 | MAAA | 7.23 | MTTC | 8.11 |
| GTMA | 7.32 | ATMA | 7.73 | CTMA | 7.71 | TTMA | 7.76 | MAAC | 7.75 | MTTT | 7.55 |
| GTMC | 7.77 | ATMC | 8.21 | CTMC | 7.94 | TTMC | 8.07 | MAAT | 7.72 | MTTM | 7.33 |
| GTMT | 7.74 | ATMT | 8.18 | CTMT | 7.81 | TTMT | 7.94 | MAAM | 7.76 | MTMG | 7.27 |
| GTMM | 7.78 | ATMM | 8.21 | CTMM | 7.30 | TTMM | 7.69 | MACG | 7.23 | MTMA | 7.70 |
| GMGG | 7.09 | AMGG | 6.93 | CMGG | 6.87 | TMGG | 6.98 | MACA | 7.65 | MTMC | 7.94 |
| GMGA | 7.34 | AMGA | 7.18 | CMGA | 7.12 | TMGA | 7.23 | MACC | 7.82 | MTMT | 7.81 |
| GMGC | 7.05 | AMGC | 7.21 | CMGC | 7.15 | TMGC | 7.26 | MACT | 7.77 | MTMM | 7.29 |
| GMGT | 7.35 | AMGT | 7.19 | CMGT | 7.13 | TMGT | 7.24 | MACM | 7.82 | MMGG | 6.86 |
| GMGM | 7.36 | AMGM | 7.20 | CMGM | 7.14 | TMGM | 7.25 | MATG | 7.54 | MMGA | 7.11 |
| GMAG | 7.42 | AMAG | 7.33 | CMAG | 7.30 | TMAG | 7.36 | MATA | 7.71 | MMGC | 7.14 |
| GMAA | 7.75 | AMAA | 7.37 | CMAA | 7.33 | TMAA | 7.40 | MATC | 7.74 | MMGT | 7.12 |
| GMAC | 7.76 | AMAC | 7.70 | CMAC | 7.64 | TMAC | 7.75 | MATT | 7.68 | MMGM | 7.14 |
| GMAT | 7.74 | AMAT | 7.63 | CMAT | 7.57 | TMAT | 7.68 | MATM | 7.74 | MMAG | 7.30 |
| GMAM | 7.76 | AMAM | 7.69 | CMAM | 7.63 | TMAM | 7.75 | MAMG | 7.21 | MMAA | 7.32 |
| GMCG | 7.28 | AMCG | 7.20 | CMCG | 7.17 | TMCG | 7.22 | MAMA | 7.64 | MMAC | 7.63 |
| GMCA | 7.71 | AMCA | 7.63 | CMCA | 7.60 | TMCA | 7.65 | MAMC | 7.82 | MMAT | 7.56 |
| GMCC | 7.79 | AMCC | 7.64 | CMCC | 7.88 | TMCC | 7.49 | MAMT | 7.76 | MMAM | 7.63 |
| GMCT | 7.77 | AMCT | 8.26 | CMCT | 7.80 | TMCT | 8.10 | MAMM | 7.82 | MMCG | 7.17 |
| GMCM | 7.27 | AMCM | 7.18 | CMCM | 7.51 | TMCM | 7.57 | MCGG | 6.87 | MMCA | 7.60 |
| GMTG | 7.60 | AMTG | 7.52 | CMTG | 7.49 | TMTG | 7.54 | MCGA | 7.13 | MMCC | 7.86 |
| GMTA | 7.72 | AMTA | 7.94 | CMTA | 7.91 | TMTA | 7.96 | MCGC | 7.15 | MMCT | 7.78 |
| GMTC | 7.73 | AMTC | 8.22 | CMTC | 7.76 | TMTC | 8.04 | MCGT | 7.14 | MMCM | 7.12 |
| GMTT | 7.70 | AMTT | 8.19 | CMTT | 7.67 | TMTT | 7.95 | MCGM | 7.15 | MMTG | 7.48 |
| GMTM | 7.73 | AMTM | 8.22 | CMTM | 7.25 | TMTM | 7.32 | MCAG | 7.30 | MMTA | 7.90 |
| GMMG | 7.27 | AMMG | 7.19 | CMMG | 7.16 | TMMG | 7.21 | MCAA | 7.33 | MMTC | 7.74 |
| GMMA | 7.70 | AMMA | 7.62 | CMMA | 7.59 | TMMA | 7.64 | MCAC | 7.65 | MMTT | 7.65 |
| GMMC | 7.55 | AMMC | 8.15 | CMMC | 7.23 | TMMC | 7.76 | MCAT | 7.58 | MMTM | 7.74 |
| GMMT | 7.77 | AMMT | 8.26 | CMMT | 7.52 | TMMT | 7.64 | MCAM | 7.64 | MMMG | 7.15 |
| GMMM | 7.26 | AMMM | 7.16 | CMMM | 7.02 | TMMM | 7.13 | MCCG | 7.17 | MMMA | 7.58 |
| | | | | | | MMMC | 7.21 | MMMT | 7.12 | MMMM | 7.02 |

Table S5. Calculated vIP values of quadruplets of successive nucleobases in B1 conformation (in eV) containing at least one 5mC (M).
